# Humans induce differential access to prey for large African carnivores

**DOI:** 10.1101/2020.06.09.143016

**Authors:** Kirby L. Mills, Nyeema C. Harris

**Affiliations:** Applied Wildlife Ecology Lab, Ecology and Evolutionary Biology Department, University of Michigan, Ann Arbor, MI 48109, USA

**Keywords:** Apex predators, diel period, encounter rates, nocturnal, prey, protected area, temporal partitioning, *Panthera leo*, West Africa

## Abstract

Wildlife adaptively respond to human presence by adjusting their temporal niche, possibly modifying encounter rates among species and trophic dynamics that structure communities. Here we show that these human-induced modifications to behaviours are prolific among species and alter apex predators’ access to prey resources. We assessed human-induced changes to wildlife diel activity and consequential changes in predator-prey overlap using 11,954 detections of three apex predators and 13 ungulates across 21,430 trap-nights in West Africa. Over two-thirds of species altered their diel use in response to human presence, and ungulate nocturnal activity increased by 6.8%. Rather than traditional pairwise predator-prey comparisons, we considered spatiotemporally explicit predator access to a suite of prey resources to evaluate community-level trophic responses to human presence. Although leopard prey access was not affected, lion and hyena access to 3 prey species significantly increased when prey increased their nocturnal temporal niche to avoid humans. Ultimately, humans considerably altered the composition of available prey, with implications for prey selection, demonstrating how humans perturb ecological processes via behavioural modifications.

---

The diel activity of wildlife can adaptively respond to their environment by partitioning time to maximize survival and limit exposure to risks, producing a species’ temporal niche^1,2^. Prey commonly employ predator avoidance strategies along the temporal niche axis^3^, which is contrasted by predators selecting for temporal activity patterns that maximize hunting success and minimize competitive encounters^4,5^. As a result, large carnivores are predominantly nocturnal while ungulates often exhibit more diurnal behaviour, though neither exclusively so. However, pervasive human pressures disrupt individual behaviours that facilitate coexistence of predator and prey populations alike^6,7^. How the human-induced responses of many species cascade to alter the dynamics of predation and other ecological interactions at the community level remains understudied.

The fear of humans suppresses spatiotemporal activity in both carnivores and herbivores with cascading impacts to lower trophic levels^8–10^. Specifically, human presence engenders shifts in diel activity patterns across guilds, altering their temporal niche to incorporate avoidance of human encounters^10^. Human activities concentrated in the day and predator activity at night reduce the availability of temporal refugia from risky encounters (either competitive or consumptive), thus constraining species’ abilities to optimize activity along the temporal niche axis^1,3^. As predator and prey species alter their diel activity to adaptively respond to human presence, predator-prey temporal overlap and resulting encounter rates are likely to be changed^11^, thus altering predator access to a suite of prey resources (Fig. 1). Such perturbations to predator-prey dynamics can have cascading impacts that alter population regulation, habitat structure, and various ecosystem processes, such as carbon storage, herbivory and seed dispersal^12–16^.

**Figure 1:**
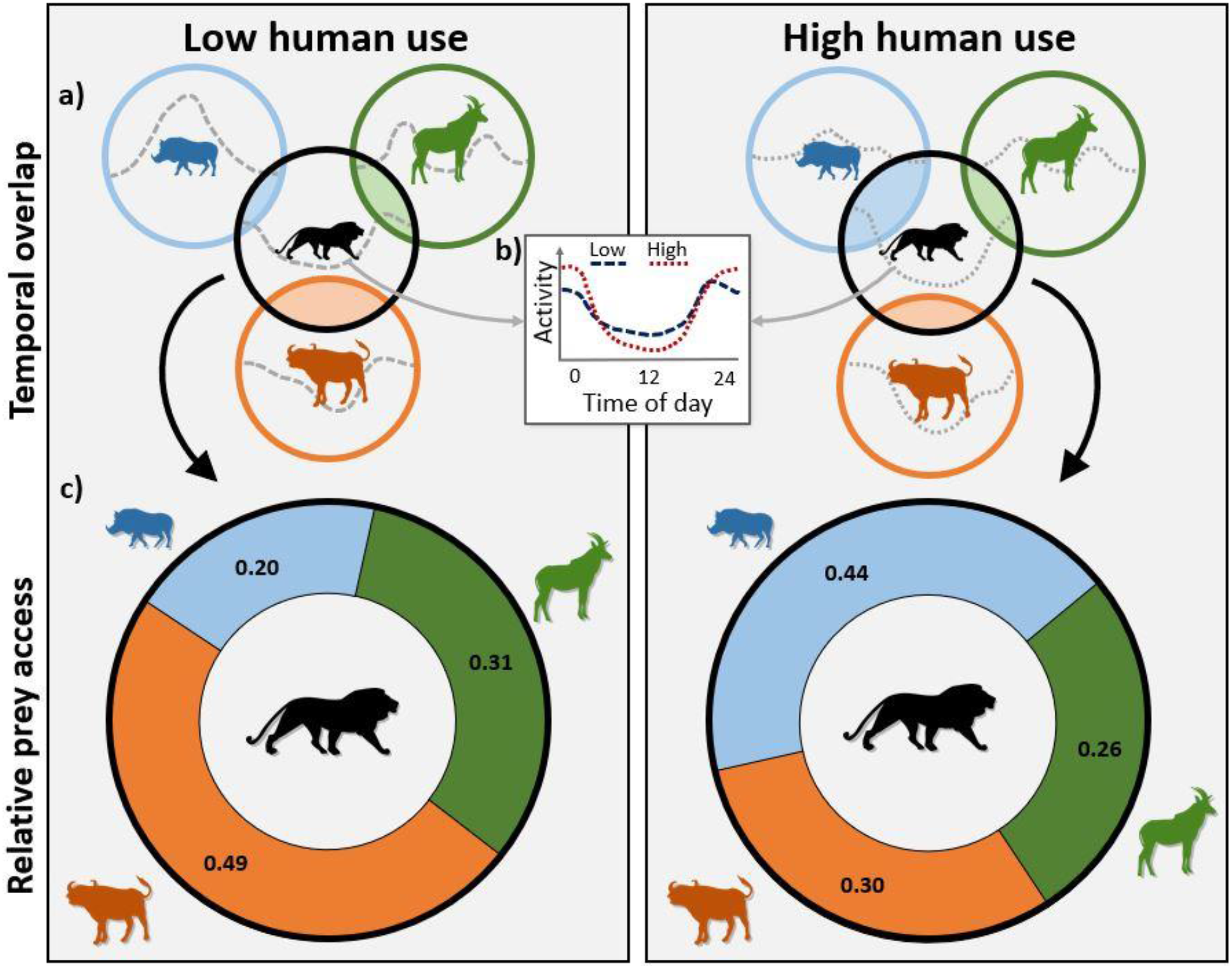
Conceptual framework illustrating the community-level effects of human presence on predator-prey temporal interactions. a) Circles represent the temporal niche occupied by each species (3 prey and 1 predator), and shaded regions indicate temporal overlap between the predator and a prey species (i.e., shared temporal niche space). Dotted lines within each circle depict the species’ temporal activity distribution. b) The diel activity patterns of both predators and prey are expected to shift in response to human presence, generally increasing nocturnal activity to avoid humans during the day. c) As wildlife diel activity changes, so does predator access to individual prey. This can lead to intensified or relaxed predation pressures on an individual prey species depending on the diel responses of the prey compared to the predators and other sympatric prey.

If wildlife modify their temporal niche to avoid pressures associated with human presence, predators and prey will exhibit increased nocturnal activity at both the species and guild levels^10^. If all species respond to humans similarly, this human avoidance hypothesis further predicts: 1) intensified predator-prey overlap overall, and 2) a greater diversity of prey species available to predators as previously diurnal species adopt nocturnal behaviours. Increasing the diversity of accessible prey would likely result in diminished predation rates on individual species, given that prey selection by carnivores is influenced by prey species’ availability relative to other sympatric prey and the diversity of the prey community^17,18^.

However, avoidance of humans may not be ubiquitous across species given species have different vulnerabilities to humans^19^. Thus, the prevalence of human avoidance among species is likely to determine the nature of community-level predator-prey outcomes.

Here, we evaluated the effects of human presence on the diel activity of predators and prey and consequential alterations to predator-prey relationships using a novel method to assess predator-prey overlap at a community scale. We executed a systematic camera survey spanning 13,100-km^2^ of the W-Arly-Pendjari (WAP) complex in West Africa across 21,430 trap-nights, obtaining detections of both wildlife and humans. We used occupancy modelling to determine areas of low and high human use within the study area and evaluate spatially explicit responses in species’ behaviour and potential alterations to trophic interactions. Specifically, we tested for shifts in diel activity patterns and nocturnal behaviours for three apex predators and 13 ungulate species between areas of low and high human presence. We also evaluated the effects of human presence on the overall temporal overlap (∆) between each predator and its prey, as well as assessed changes in the relative overlap between predators and each individual prey species. We determined: i) how carnivores and ungulates adjusted their temporal niche in response to human presence, and ii) how apex predator access to prey species was altered by human presence.

Previous works often investigate temporal overlap of predators and prey in a pairwise manner^11,20,21^. However, such an approach does not consider the overall composition of resources available to predators and the relative contributions of individual prey species. Higher-order interactions beyond pairwise predator-prey relationships likely contribute to determining community structure and coexistence among species^22^. We combatted these limitations by extending beyond pairwise comparisons to consider predator-prey interactions at the community level by aggregating temporal activity among ungulates, providing a more ecologically realistic depiction of overlap between predators and their prey.

Specifically, we used bootstrapped kernel density distributions of predator and prey diel activity to calculate the overlap between each predator-prey pair relative to the overall available prey (percent area under the predator diel curve, PAUC), which was generated by aggregating prey activity curves and then scaling the prey activity (kernel density estimates) to each predator. PAUC represents a metric of relative prey access for the apex predator, as it provides insight into the times of day that encounters between the predator and prey species are most likely to occur based on the temporal activity of both. In assessing the spatially explicit temporal responses of predators and their prey to humans, we elucidate the community-level effects of humans on trophic interactions and their implications for ecosystem regulation by large carnivores.

## RESULTS

Our camera survey yielded 786 and 11,168 detections of apex predators and ungulates, respectively, over 21,430 trap-nights throughout our West African study system (Table S1). Hyenas are the dominant predator in the system with 6 times more detections than either lions or leopards. Warthog, reedbuck, and bushbuck were the most commonly observed ungulates, each detected over 1,000 times.

We obtained 704 detections of humans in 69 out 204 surveyed 10-km^2^ grid cells, leading to a naïve human occupancy of 0.34. Accounting for imperfect detection, model selection resulted in four competing top models (∆AICc < 2) for human occupancy (Table S2). Detection of humans primarily varied among years and sites and was higher in non-savanna habitat (top model goodness-of-fit *p-value* = 0.327).

Human occupancy was pervasive, but heterogeneous within the study area (Fig. 2a; 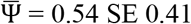), ranging from 0.0006 to 1 with highest frequencies near these extremes (Fig. 2b). Using the mean value of occupancy as a pressure threshold, we designated 108 of 204 grid cells as having high human use (occupancy > 0.54).

**Figure. 2:**
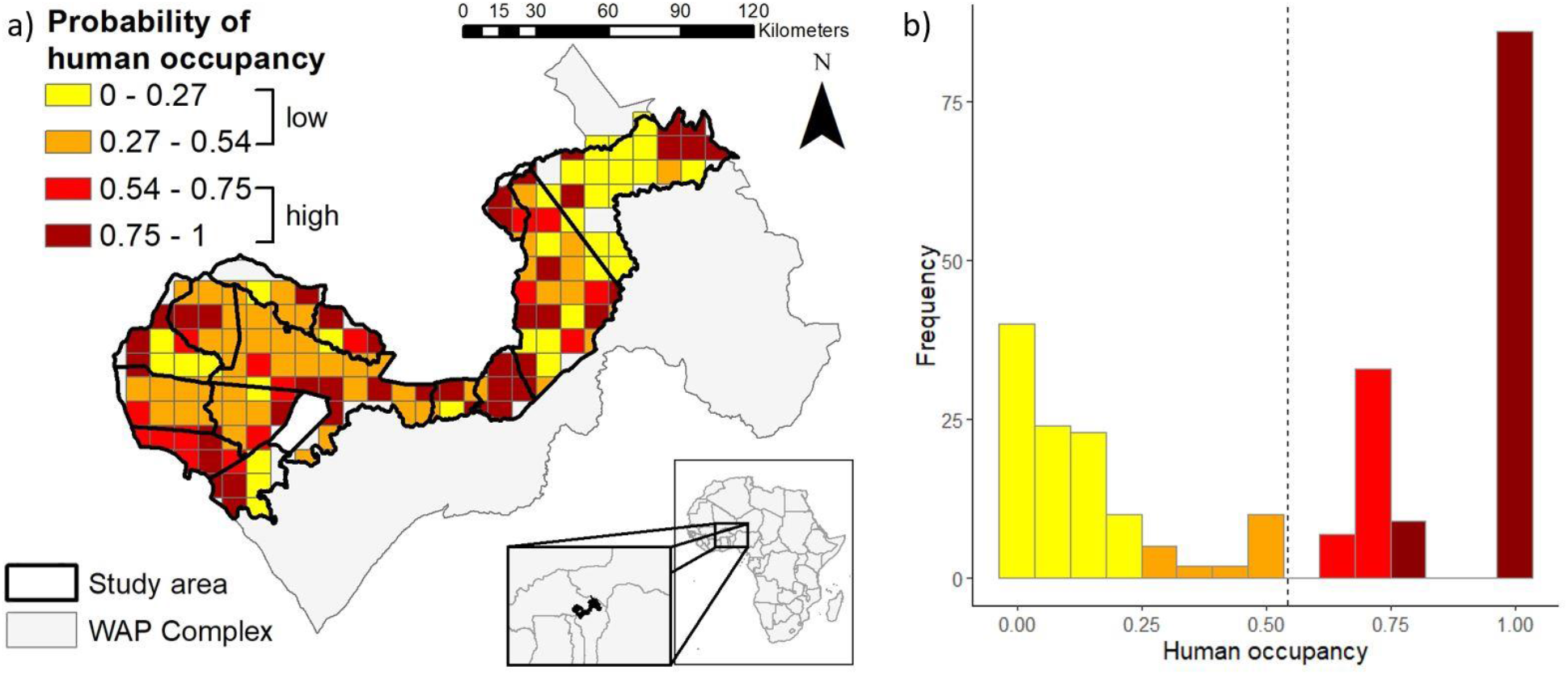
a) Map of study area within the W-Arly-Pendjari protected area complex in Burkina Faso and Benin, West Africa. Surveyed 10-km^2^ grid cells are depicted showing estimated levels of human occupancy within the study area, averaged across years for grid cells surveyed in multiple years. b) Corresponding frequencies of grid-level human occupancy for 3 survey years, with dotted line depicting mean human occupancy (0.54).

### Human avoidance responses

Human presence generated marked modifications in the temporal niches of sympatric wildlife, with both guilds exhibiting human avoidance behaviours overall. Carnivores and ungulates significantly altered their diel activity under high human presence (carnivores *p-value* = 0.017; ungulates *p-value* < 0.001; Fig. 3). Over two-thirds (11 out of 16) of the mammal species in the study exhibited significant shifts in their diel activity patterns in response to human presence (leopards, hyenas, and 9 ungulates; Fig. 3, Table S3). Ungulates overall became 6.7% (95% CI ± 1.5%) more active at night in high human areas, while carnivores showed a slight but non-significant increase in night-time activity of 3.9% (± 5.7%).

**Figure 3:**
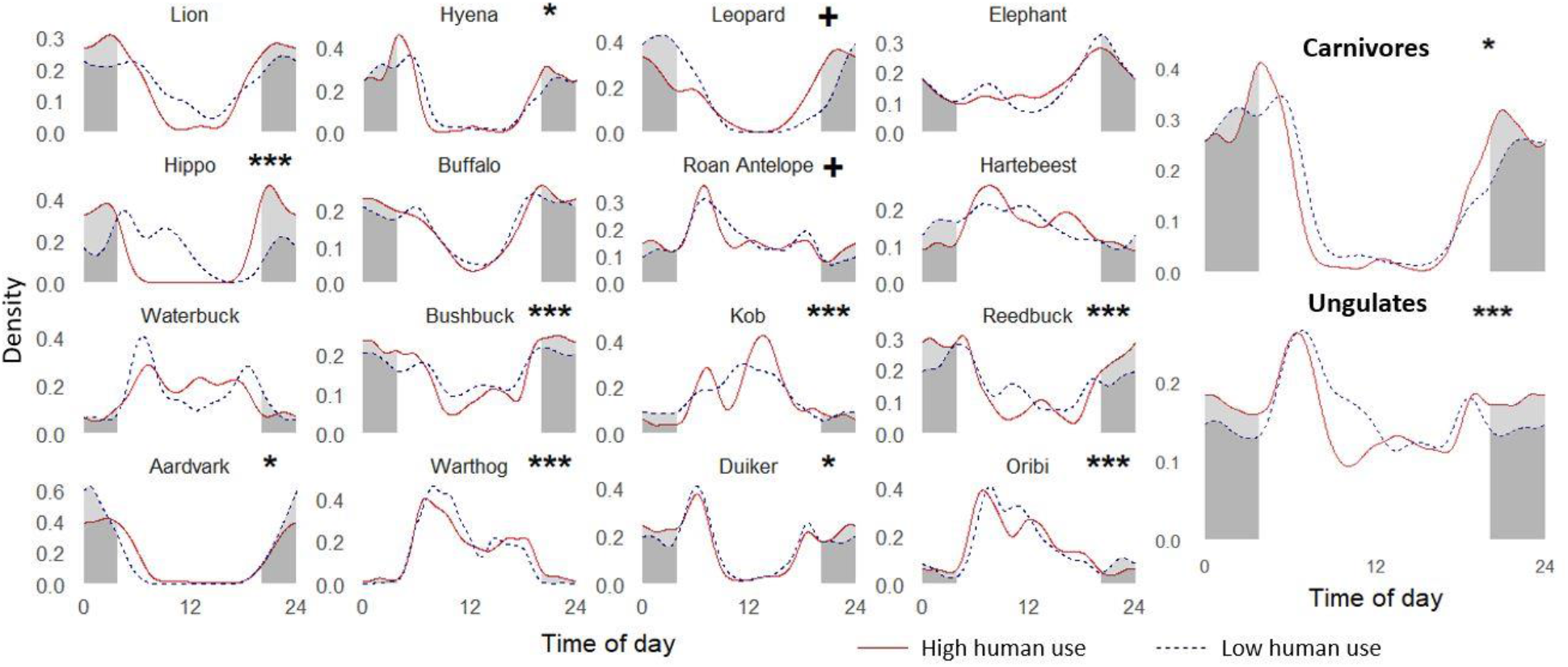
Temporal activity kernel density curves for apex predators and ungulates in areas of low and high human use (threshold human occupancy = 0.54). Nocturnal diel periods (2 hours after sunset to 2 hours before sunrise) are shaded using the average times of sunrise and sunset during our study period, and lighter shading represents the diel-specific nocturnal activity that is different between low and high human areas. Significance levels for bootstrapped randomization test of differences in diel distributions between human zones: * < 0.05, ** < 0.01, *** < 0.001. Plus signs (**+**) represent species with p-values < 0.1 which achieved significance when the human occupancy threshold was adjusted ± 0.1 (Table S3).

Specifically, we observed significant increases in nocturnal activity with human presence for hippopotamus (+35.0 ± 18.0%), reedbuck (+12.3 ± 4.8%), duiker (+7.4 ± 4.4%), bushbuck (+6.9 ± 3.6%), and warthog (+4.5 ± 1.9%); and significant decreases for kob (−5.3 ± 4.2%) and aardvark (−15.0 ± 8.1%; Fig. 4). In contrast, 6 ungulate species and all 3 carnivores showed no significant changes in nocturnality.

**Figure 4:**
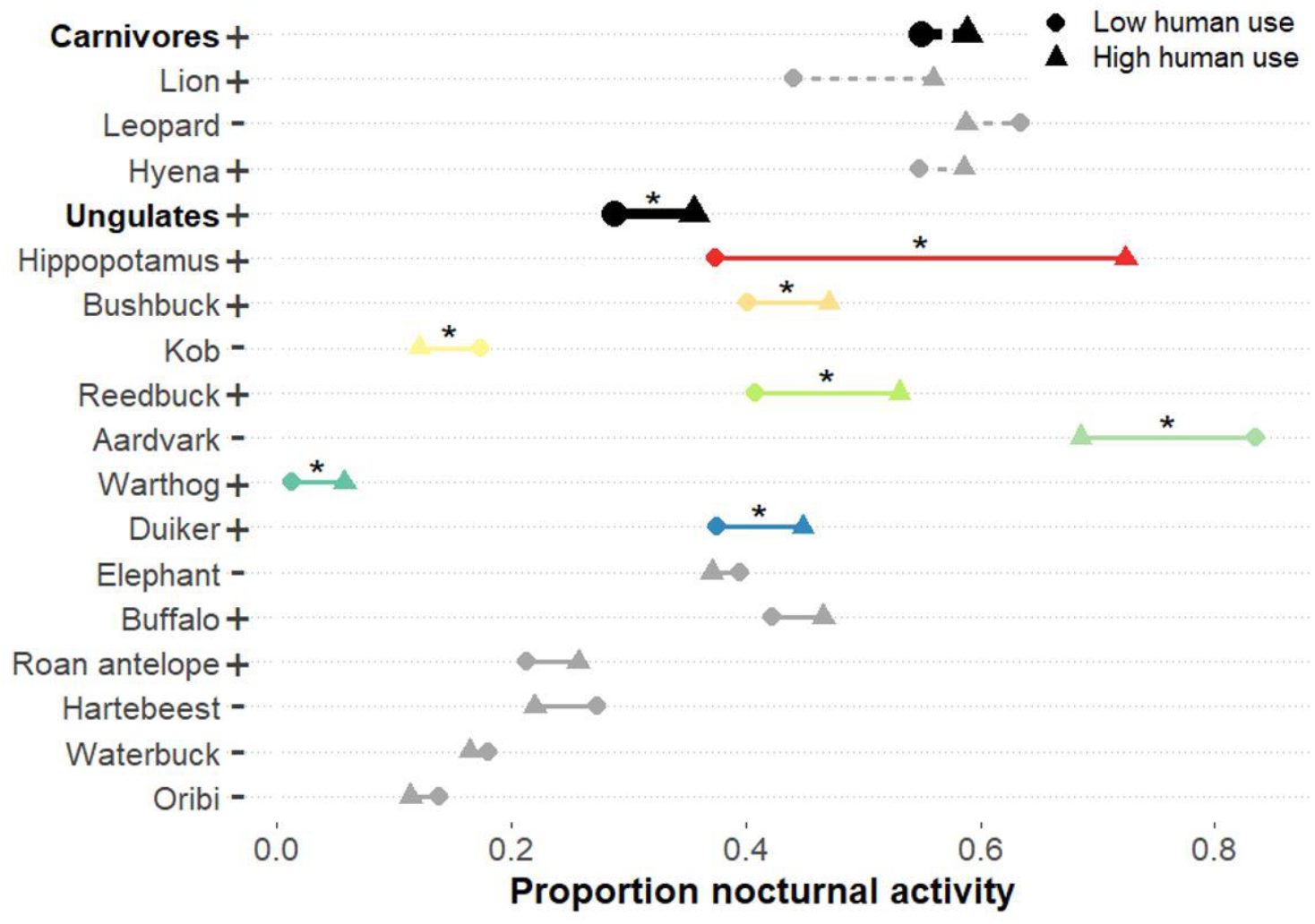
Proportion of activity during the nocturnal diel period (2 hours after sunset to 2 hours before sunrise) between low and high human zones for carnivore (dashed lines) and ungulate (solid lines) species and overall guilds. Increases and decreases in nocturnality from low to high human use areas are indicated by plus (+) and minus (−) labels, respectively, next to species’ names. Stars (*) above colored lines indicate species that showed significant differences in nocturnal activity between human zones based on bootstrapped 95% CIs of nocturnality.

Overall, our threshold of human occupancy designating low vs. high pressure did not alter our interpretation of sensitivity from resultant shifts in diel activity. The only change we observed was detecting significance when reducing the threshold from 0.54 to 0.44 for 2 species: leopard and roan antelope (Table S3). Our results highlight that most species respond to human occurrence by modifying their behaviours and reducing their realized temporal niche to incorporate more night-time activity, potentially altering predator-prey encounter rates.

### Changes in predator-prey overlap

Human-induced diel activity shifts among species did not result in significant changes in individual predators’ overlap with prey when we aggregated their prey species (Fig. S2). However, high human use reduced the mean overlap of lions with their prey by 0.08 (∆_high_ = 0.718 ± 95% CI 0.08; ∆_low_ = 0.797 ± 0.11). In contrast, leopards may be experiencing some benefit from human use, as their temporal overlap with prey exhibited a mean increase of 0.17 where human activities were high (∆_high_ = 0.699 ± 0.12; ∆_low_ = 0.529 ± 0.11). Hyenas appear to be robust to human occurrence, showing almost no changes in total overlap with prey due to humans (∆_high_ = 0.638 ± 0.03; ∆_low_ = 0.625 ± 0.04).

### Human occurrence restructures access to specific prey

Lions and hyenas similarly experienced distinct changes in the composition of accessible prey due to human presence, evaluated from 95% CIs of ∆PAUC overlap with 0. Specifically, humans generated significant changes in overlap of these predators with 4 out of 11 prey species: bushbuck (∆PAUC_lion_ = +1.49 ± 95% CI 1.14%, ∆PAUC_hyena_ = +1.51 ± 0.73%), reedbuck (∆PAUC_lion_ = +1.99 ± 1.34%, ∆PAUC_hyena_ = +1.88 ± 0.87%), duiker (∆PAUC_lion_ = +1.56 ± 1.31%, ∆PAUC_hyena_ = +0.84 ± 0.73%), and kob (∆PAUC_lion_ = −2.24 ± 1.58%, ∆PAUC_hyena_ = −1.45 ± 0.88%) (Fig. 5). All three species to which predator access increased significantly also exhibited increased night-time activity as a human avoidance strategy. In contrast, kob reduced nocturnality and experienced lower overlap with lions and hyenas (Fig. 4). Additionally, lion and hyena access to 2 prey species showed near significant changes (buffalo ∆PAUC_lion_ = +1.33 ± 1.41%, ∆PAUC_hyena_ = +0.87 ± 0.97%; and waterbuck ∆PAUC_lion_ = −1.66 ± 1.73%, ∆PAUC_hyena_ = −1.67 ± 1.71%).

**Figure 5:**
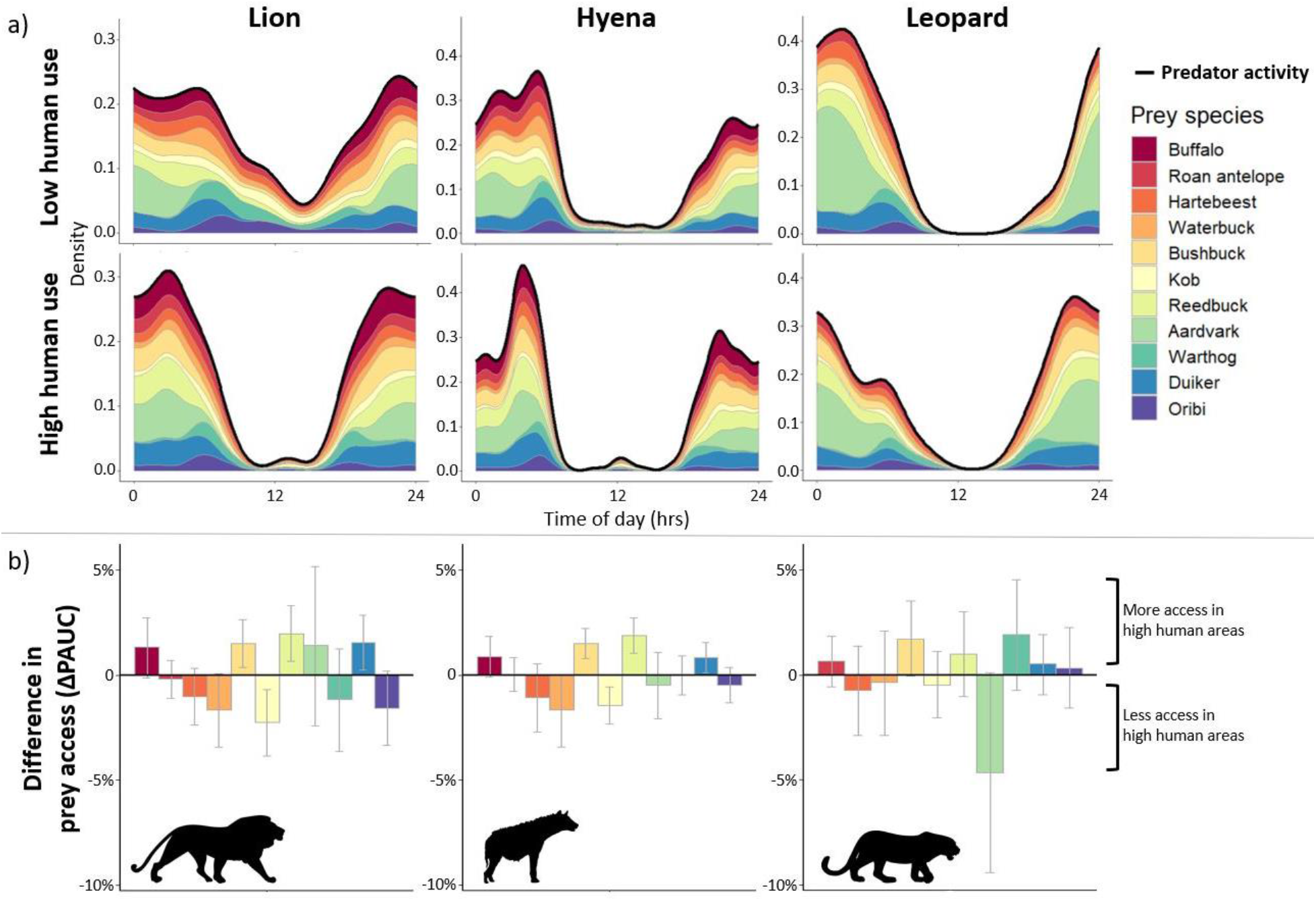
Differences in prey access between human activity zones for lions, leopards, and hyenas. Buffalo was not included as prey for leopard. a) Temporal overlap heatmaps representing the relative contributions of each prey species to the overall available prey base scaled to predator activity curve (solid black line) over the 24-hour cycle. Density values are calculated from kernel density temporal curves for predators and prey species. b) Mean differences in prey access based on species-specific area under the predator activity curve (∆PAUC) between areas of low and high human pressure, averaged among 10,000 parametric bootstrap replicates. Error bars represent bootstrapped 95% confidence intervals of ∆PAUC. Positive values of ∆PAUC indicate more access to a prey species in high human use areas relative to low, while negative values indicate less access in high human use areas.

All three apex predators showed strong similarities in prey access differences with comparable changes in access to all prey between human use levels (Fig. 5). However, changes in leopard access to prey items were not significant based on 95% CIs, with only aardvark (∆PAUC_leopard_ = −4.6 ± 4.7%) and bushbuck (∆PAUC_leopard_ = 1.7 ± 1.8%) access nearing significance (Fig. 5). We suspect this is due to leopards’ differential response to human presence (−4.6 ± 19.1% change in nocturnality) compared to lions (+11.9 ± 16.3%) and hyenas (+3.7 ± 6.5%; Fig. 4).

Although overlap with total available prey did not differ, the variation in species-specific prey accessibility (PAUC estimates) was higher for lions as a result of human presence (Fligner-Killeen test, *p-value* = 0.03), indicating reduced diversity of available prey and therefore increased access to certain prey species compared to others. Leopards and hyenas showed no significant changes in access variability as a response to humans.

## Discussion

Wildlife responses to human activities have the potential to reshape natural ecological processes and trophic dynamics^8,9,23^. When anthropogenic pressures are heterogeneous, the resultant dynamism promotes a myriad of adaptive strategies to manage and mitigate threats including behavioural shifts in diel activity that redefine species’ temporal niches^10,24,25^. Such shifts in diel activity may lead to increased prey vulnerability to nocturnal predators, thus altering probabilities of encounter and diets in consumers (Fig. 1). We found that over two-thirds of the assessed species altered their overall diel activity patterns as a response to humans in the study area. Most species increased nocturnal activity, consistent with previous works and supporting our hypothesis of human avoidance^10,25^. Valeix *et al.*^26^ and Suraci *et al.*^27^ similarly found reduced diurnal activity near human settlements in African lions in Makgadikgadi Pans National Park, Botswana, and Laikipia, Kenya, respectively, likely to reduce risks of human encounters. Human presence appears to be limiting temporal refugia from risks for many species and driving increases in ungulate activity when predators are also active, possibly decoupling anti-predator behaviours from predation risks^28,29^. Patten *et al.*^11^ also presented evidence of human avoidance driving increased predation risks in North American white-tailed deer (*Odocoileus virginianus*).

Heterogeneity in species’ responses to human pressure, however, indicates different sensitivities to humans among the carnivores and ungulates in our study system. Some species did not exhibit shifts towards nocturnality as expected (e.g., kob and aardvark). These species may be benefitting from the observed human avoidance in many sympatric species that potentially reduces risks of predation and competition, commonly referred to as a human shield response^24,30^. For example, Atickem *et al.*^31^ reported mountain nyala (*Tragelaphus buxtoni*) leveraging predator avoidance of humans during the day as a temporal refuge in Ethiopia. The ability to exploit human presence as a shield from predatory or competitive encounters may be due to the life history traits of a species that reduce sensitivity to humans, such as body size, energetic requirements, dispersal abilities, social structure, or foraging strategies^19,32^. Similarly, these species’ temporal niches may be constrained by inherent characteristics that were evolved for diurnal activity, making night-time activity more costly despite refuge from human pressures and limiting their adaptive capacity to avoid humans^33^. In contrast, the amount of wildlife persecution (i.e., trophy hunting and poaching) in the system may induce stronger human-avoidance behaviours in hunted species. For instance, Vanthomme *et al.*^34^ attributed the negative associations of 10 mammals with human disturbances in Gabon to hunting avoidance behaviours, contrasted by 6 species in the study showing positive associations. Though the mechanisms driving differential responses to humans were not explicitly investigated here, our study demonstrates non-uniform responses of large mammals to human presence. As such, future work can assess the drivers of species-specific responses and sensitivities to humans.

We showed that human presence modified the availability of prey species relative to the overall pool of available prey, which is an important driver of prey selection in apex predators, and thus provide new insights into community-level repercussions of human sympatry with wildlife^17,18^. While we expected overall predator-prey overlap and the diversity of available prey to increase due to human avoidance, the combination of human avoidance and human shield strategies observed in our system resulted in little change in overall overlap but substantial changes to apex predator access to individual prey species. Specifically, our new community-level approach to predator-prey temporal overlap revealed that prey species experienced intensified overlap with predators when they increased their nocturnal temporal niche (e.g., duiker, reedbuck, bushbuck) to avoid humans, while overlap was lessened for species that did not (e.g., kob). For lions, this resulted in reduced diversity of available prey, likely intensifying predation pressures on a smaller subset of species which could contribute to destabilizing trophic dynamics^35^. The predators in our study are largely opportunistic night-time hunters, and temporal overlap is often strongest between predators and their preferred prey species^20,21,36,37^. Thus, we expect that species experiencing the highest overlap with apex predators relative to other prey to be integrated into the predators’ diets in higher proportions, and consequently expect varied prey selection by predators between low and high human use areas. Buffalo are a common prey item of lions in other systems, and our results suggest they may be vulnerable to intensified selection by lions due to human presence increasing access to buffalo in our study area^38^. Additionally, human disturbance can increase the predation rates and carcass abandonment by large carnivores as well as alter mesopredator foraging behaviours, potentially increasing mortality rates on preferred prey species and providing augmented carrion resources that may be detrimental to scavenger populations ^39,40^. As such, disturbances to predator-prey relationships potentially lead to alterations in predators’ diets with consequences for ungulate and mesopredator community regulation^41^.

Though protected areas are the primary strategy for biodiversity conservation worldwide, human exploitation of protected areas is pervasive and in many cases necessary for the sustenance of human populations^42,43^. By accounting for imperfect detection to understand human intensity of use, we contribute to a more comprehensive understanding of human impacts within coupled human-natural ecosystems that is imperative to effectively manage for the conservation of ecological processes, biodiversity, and human needs. However, human activities observed in our study system may not impact species uniformly. Because we aggregated a variety of human activities to depict human use, there might be activity-specific responses by wildlife that were not captured. Humans exploit resources in national parks in many ways including livestock herding, resource gathering, subsistence poaching, hunting, and recreation, all of which impact the system and wildlife to varying degrees^43–45^. Overall, human impacts encompass a variety of disturbances that impact ecosystems, both in our study and more broadly, and thus disentangling the responses of wildlife to specific human pressures may facilitate designing more effective conservation interventions^42,46^.

Our results demonstrate prevalent disruptions to wildlife temporal activity patterns from human presence, leading to overall reductions in diurnal activity and modified community dynamics. Because both carnivores and ungulates serve fundamental roles in regulating African ecosystems via predation and herbivory, respectively, the pervasiveness of their responses to human occurrence demonstrates the capacity for humans to disrupt essential ecological processes that facilitate coexistence among wildlife, in this case reshaping predator-prey interactions. As the human footprint continually expands, spatial refugia from anthropogenic disturbance become more limited, stimulating an increasing need to exploit temporal partitioning to avoid human pressures. We show that the community-level implications of these behavioural modifications must be considered in light of complex higher-order interactions that govern mechanisms of coexistence among predators and their prey.

## METHODS

### Study area

We conducted our study in the W-Arly-Pendjari (WAP) protected area complex that spans 26,515-km^2^ in the transboundary region of Burkina Faso, Niger, and Benin (0°E-3° E, 10°N-13°N; Fig. 2a). The complex contains 5 national parks (54% of total area), 14 hunting concessions (40%), and 1 faunal reserve (6%). Our study area within WAP comprised three national parks and 11 hunting concessions in Burkina Faso and Niger across ca. 13,100-km^2^ (Fig. 2a). Trophy hunting of many ungulate species and African lions (*Panthera leo*) is permitted in hunting concessions, while all hunting is illegal in the national parks and reserves in the complex. Other human activities in the park include livestock herding, resource extraction, recreation, and poaching ^44,47,48^. Recently, Harris et al. ^44^ reported 4 large carnivore species (lion, leopard *Panthera pardus*, and spotted hyena *Crocuta crocuta,* cheetah *Acinonyx jubatus*) and 17 ungulate species belonging to the superorder Ungulata in the three national parks included in our study area from an extensive camera trap survey. Cheetahs were detected only once, while wild dogs (*Lycaon pictus*) were not reported in the survey area. WAP has an arid climate and consists predominantly of Sudanian and Sahel savannas ^49^. We conducted our survey in the drier northern portion of WAP during the dry season with average monthly rainfall ranging from 0-1 mm in February to 42-91 mm in June ^50^.

### Camera survey

We systematically deployed 238 white-flash and infrared motion-sensor cameras (Reconyx© PC800, PC850, PC900) within 10×10-km grid cells across our study area to assess effects on human presence of diel activity within the wildlife community. A single unbaited camera was placed within 2-km of the centroid in a total of 204 sampled grid cells over 3 survey seasons from January-June in 2016-2018 (Fig. S1). Species identifications from camera images were validated by two members of the Applied Wildlife Ecology (AWE) Lab at the University of Michigan. We excluded false triggers, unidentifiable images, research team, and park staff from analyses. To ensure robustness in our analyses, we combined all remaining human images into a single ‘Human’ categorization representing a variety of human activities observed in WAP (e.g., livestock herding, resource gathering, recreation, poaching, and hunting). (see Fig. S1, Mills *et al.* ^51^ and Harris *et al.* ^44^ for additional methods on camera deployment and image processing). We created independence of species triggers using a 30-minute quiet period between detection events using the ‘camtrapR’ package in R 3.5.1 (http://www.r-project.org) ^52^, and we assumed detections to be a random sample of each species’ underlying activity distribution ^21^.

### Human occupancy models

We constructed single-season, single-species occupancy models to discriminate WAP into areas of low and high human use. We separated detection/non-detection data for humans into 2-week observation periods, which were modelled as independent surveys to account for imperfect detection. Our occupancy models first modelled the detection process (*p*) using covariates expected to influence detection while holding occupancy (Ψ) constant, and then modelled human occupancy by incorporating grouping variables among which Ψ may vary.

The global detection model included covariates related to survey design and the environment that we expected to influence the detection of humans: % savanna habitat (SAV), survey year (YR), trap-nights (TN), camera type (CAM), management type (MGMT), and site (i.e., one of 14 individual parks or concessions; SITE). MGMT was a binary variable that distinguished national parks from hunting concessions. Human occupancy was modelled with only grouping variables: MGMT, YR, and SITE. We included YR as a covariate to account for temporal variation in site use or detection, as cells surveyed in multiple years were disaggregated as separate sites for our single-season model. Variables included in the top performing occupancy and detection model(s) are considered those which best described the spatial variation in human detection and site use. We evaluated the support for all combinations of detection and occupancy covariates using the Akaike information criterion corrected for small sample sizes (AICc). We selected the top-performing detection and occupancy models as those with ∆AICc < 2 compared to the lowest AICc model. We assessed goodness-of-fit of the top-performing models using 1,000 parametric bootstraps of a χ^2^ test statistic appropriate for binary data and estimated the *ĉ* statistic to ensure the data were not over-dispersed ^53^. We created all detection and occupancy models using the ‘unmarked’ package and conducted model selection using the ‘MuMIn’ package in R.

We extracted cell-specific latent occupancy probabilities, representing intensity of site use by humans because the 10-km^2^ grid cells do not meet the assumption of closure, from the top-performing (lowest AICc) occupancy model corrected for imperfect detection ^54^. From those estimates, we categorized grid cells as either low or high human use. We delineated the threshold for human use using the mean value of human occupancy, which presented a bimodal distribution at low and high levels of human site use (Fig. 2b).

### Temporal analyses

Using detection timestamps from our camera survey, we compared the temporal activity patterns for apex predators (lions, leopards, and hyenas) and sympatric ungulates between areas of low and high human use. We included 14 ungulate species: elephant (*Loxodonta africana*), hippopotamus (*Hippopotamus amphibius*), savanna buffalo (*Syncerus caffer brachyceros*), roan antelope (*Hippotragus equinus koba*), western hartebeest (*Alcelaphus buselaphus major*), waterbuck (*Kobus ellipsiprymnus defassa*), Buffon’s kob (*Kobus kob kob*), Bohor reedbuck (*Redunca redunca*), bushbuck (*Tragelaphus sylvaticus*), aardvark (*Orycteropus afer*), warthog (*Phacochoerus africanus*), oribi (*Ourebia ourebi*), red-flanked duiker (*Cephalophus rufilatus),* and common duiker (*Sylvicapra grimmia*). Due to few detections, we excluded topi (*Damaliscus korrigum jimela*) and red-fronted gazelle (*Eudorcas rufifrons*) from analysis in our study. Duiker species were aggregated due to difficulty distinguishing the two in camera trap images, resulting in 13 total ungulate species in our analyses.

We used kernel density estimation to produce diel activity curves representing a species’ realized temporal niche in both human use zones for each of the 16 species. We first tested for differences in these activity distributions between low and high human use areas for all individual species and overall for each guild by calculating the probability that two sets of circular observations come from the same distribution with a bootstrapped randomization test ^55^. Significant differences in temporal activities were evaluated as *p-value* < 0.05. We conducted a sensitivity analysis by adjusting the human occupancy threshold ± 0.1 and repeating this test for all species and both guilds to ensure robustness of our results (Table S3).

Using 10,000 parametric bootstraps of the temporal distribution models, we then calculated the area under the diel activity curves to determine the proportion of each species’ activity that occurred during nocturnal hours (two hours after sunset to two hours before sunrise). We used the sunrise (05:51) and sunset (18:06) times from the median date of our surveys (April 4, 2018) at the survey area centroid to define nocturnal hours. To test if wildlife nocturnality increased to avoid human presence, we compared the bootstrapped 95% confidence intervals (CIs) of the change in nocturnality for each species and overall guilds between low and high human areas where a significant change was observed when the CI did not overlap 0.

We used the coefficient of overlap (∆) to quantify the total temporal overlap between each apex predator and their associated prey from circular activity distributions. Elephant and hippopotamus were not included in the prey list for any predator and buffalo was excluded from the prey list of leopards due to large body sizes. All other prey species were aggregated to produce a single diel activity curve of all prey for comparison to predator activity. We chose the specific estimator based on the minimum sample size of detections for both guilds to contrast human use levels (∆1 if N < 75, ∆4 if N > 75). Values of ∆ range from 0-1 where 0 represents no temporal overlap and 1 represents complete overlap or identical temporal niche between predators and their prey. We used 10,000 bootstrapped estimates to extract the bias-corrected 95% CIs of ∆. We compared CIs of ∆ between human use levels for each species to assess differences in predator-prey overlap in response to human occurrence. Non-overlapping CIs between human use levels indicated that the overall temporal overlap of predators with their prey was significantly altered by human presence. Temporal analyses were conducted using the ‘activity’ and ‘overlap’ packages in R.

### Predator access to prey

After determining overlap between predators and their prey as well as shifts induced by humans, we determined the implications for predator access to prey. To our knowledge, we developed a new method to assess species-specific prey access for predators that is temporally explicit over the diel period, enabling assessment of changes to the composition and diversity of accessible prey for predators resulting from responses to humans in both guilds. We first combined (i.e., stacked) the bootstrapped temporal kernel density curves for individual prey to produce a total diel activity curve for prey, but this time maintaining each species’ contributions to overall prey activity. We then multiplied each prey species’ proportional contribution to prey activity at a given point in the diel cycle by the corresponding kernel density activity value of each respective apex predator. This method produced a discrete area under the predator temporal activity curve for each prey species of a given apex predator (percent area under curve, PAUC), where each prey species’ value represents the relative temporal overlap between the apex predator and that prey species throughout the day. We used these PAUC values to assess whether predator access to individual prey species, relative to all available prey, was altered by human presence by calculating the difference in prey access (∆PAUC) for each predator/prey combination between areas of low and high human use. To determine if prey access was significantly changed by human presence, we compared bootstrapped estimates and 95% CIs of ∆PAUC. Finally, we used a Fligner-Killeen test for homogeneity of variance to determine how the diversity of each predator’s accessible prey changed in association with human presence based on PAUC values. Lower variance in prey access represents more evenness (i.e., more diversity) in access across prey items, while higher variance indicates prey access is higher for a subset of species compared to others.

## Acknowledgements

We thank the AWE Lab for comments on manuscript drafts, assistance with image identification, and data management specifically S. Gamez, M. Lyons, J. VanZoeren, and R. Malhotra. We also thank B. Dantzer and J. Suraci for their valuable comments on the manuscript. We extend our sincerest appreciation to all park managers and the field team, especially I. Gnoumou and Y.I. Abdel-Nasser that assisted with data collection. We also thank administrators in Ministries of Environment in Burkina Faso (OFINAP & DGEF) and Niger (DGE/EF) especially B. Doamba and Y. Harissou for logistical support as well as private concessionaires in Burkina Faso for wildlife management efforts and access to properties in WAP. We also acknowledge the University of Michigan (UM) African Studies Center - STEM initiative, UM Office of Research, the German Society of Mammalian Biology, and the Detroit Zoological Society for financial support.

## Supplemental Information

**Humans disrupt predator-prey dynamics by inducing shifts in species’ temporal niches**

Mills and Harris

---

**Fig. S1:**
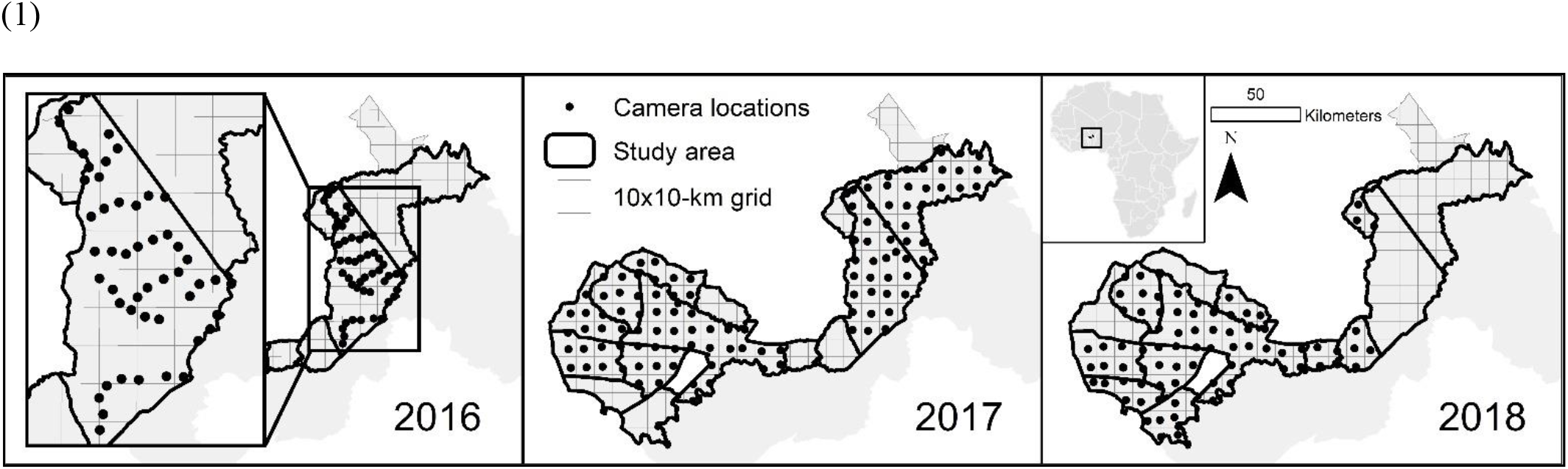
Camera placement in W-Arly-Pendjari protected area complex from three survey years. 50 cameras were deployed in 2016, 115 cameras in 2017, and 73 cameras in 2018. Modified from Mills *et al.*

**Figure S2:**
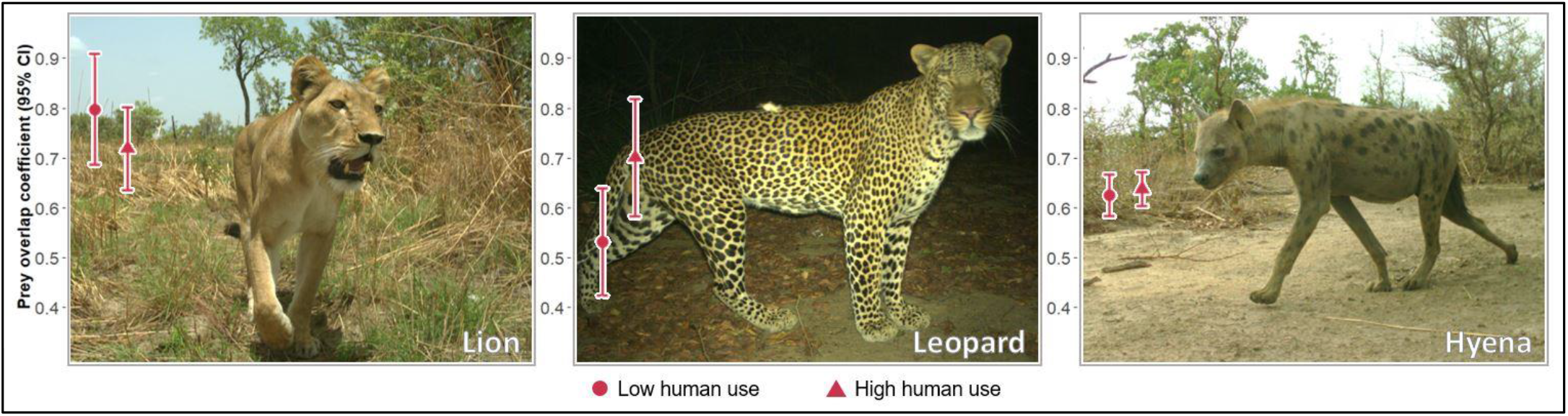
Temporal overlap coefficients (∆) between each predator and their associated prey species. Buffalo not included as prey for leopard, and elephant and hippopotamus not included for any predator. Error bars represent bias-corrected, bootstrapped 95% confidence intervals of coefficient estimates. Photo credit: Applied Wildlife Ecology Lab (AWE), University of Michigan, images from camera trap survey within the study area.

**Table S1:**
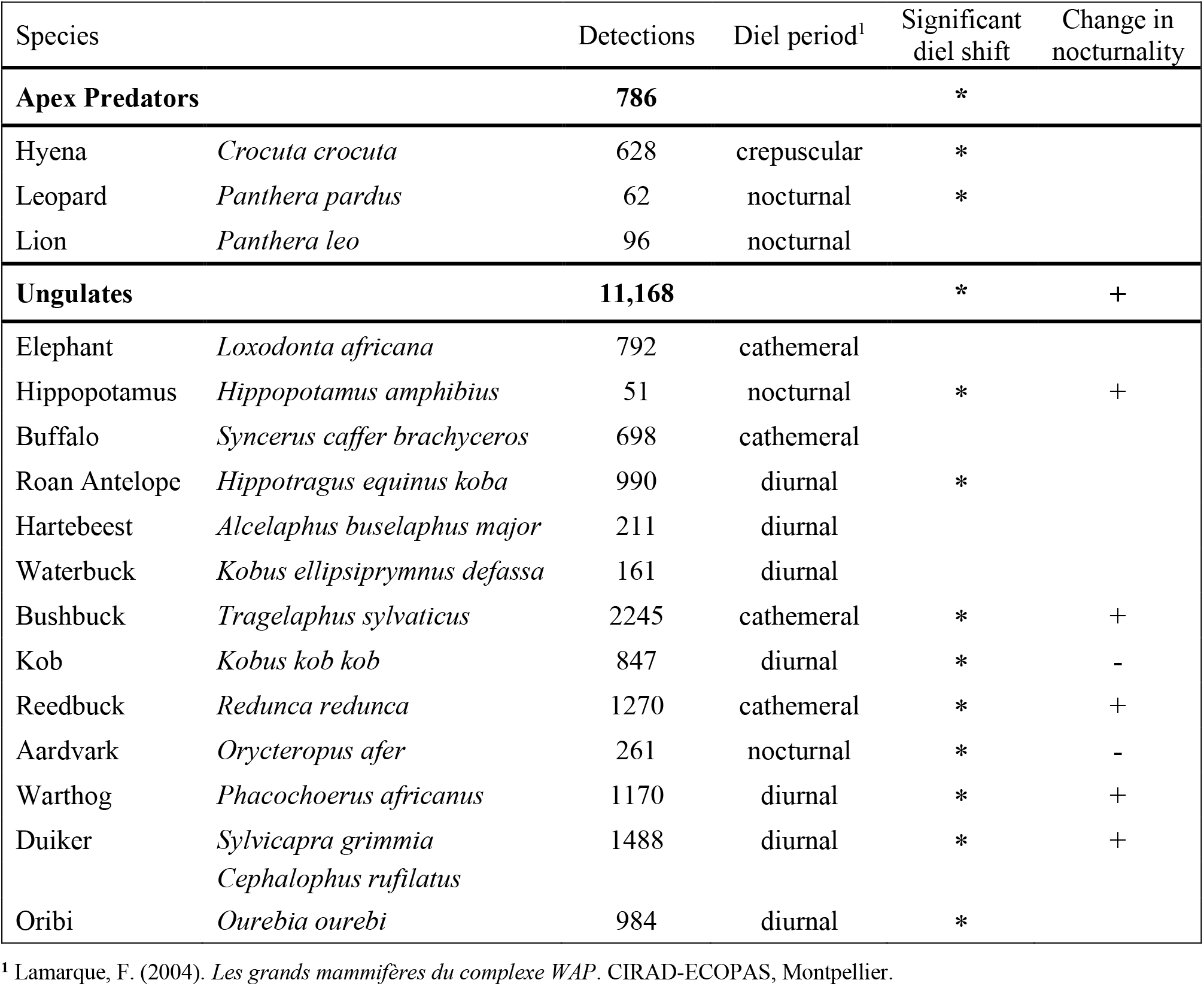
Species detections during the camera survey and common diel period. Asterisks (*) indicate significant shifts in diel activity distributions due to human presence. Changes in nocturnality are depicted for species with significant increases (+) and decreases (−) in response to humans. Empty cells represent no significant changes.

**Table S2:**
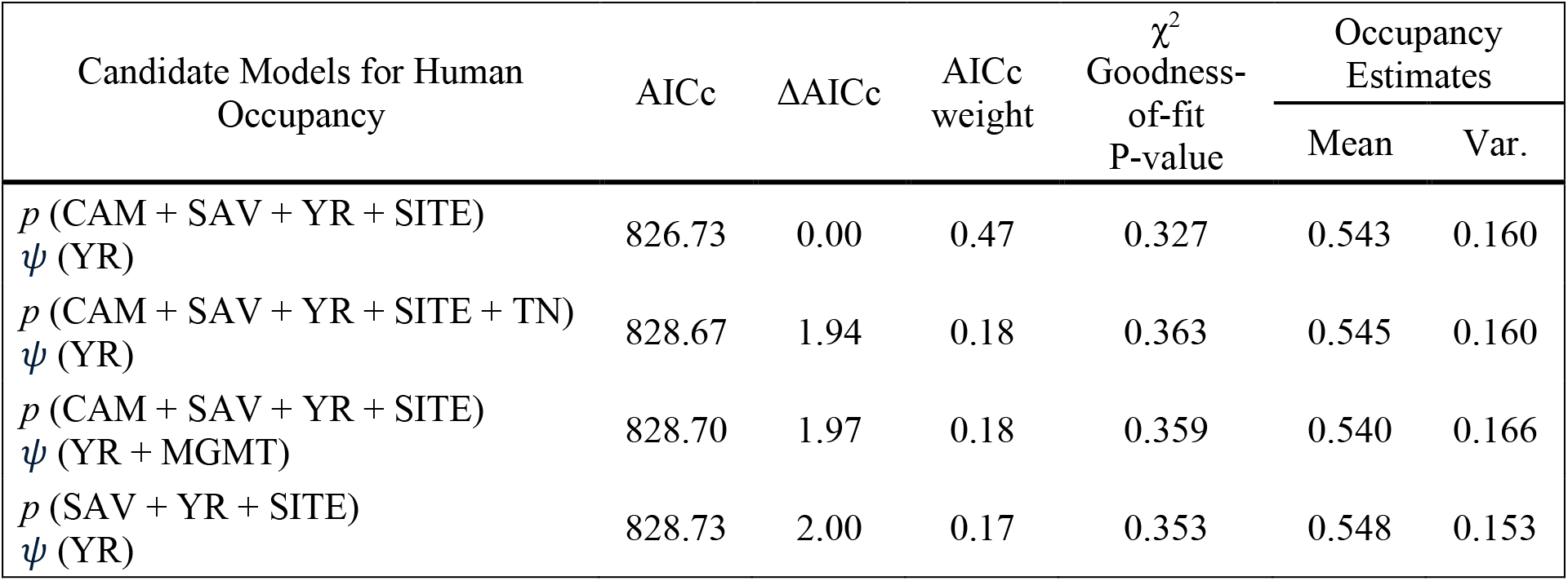
Human occupancy model selection table of top models with ∆AICc < 2 derived from camera data collected over 3 survey seasons in the W-Arly-Pendjari complex, West Africa. Detection (*p*) and occupancy (*Ψ*) were modeled using the following covariates: CAM = camera type, SAV = percent savanna, YR = survey year, SITE = survey site, MGMT = management type (national park or hunting concession).

| Candidate Models for Human Occupancy | AICc | $\Delta\text{AICc}$ | AICc weight | $\chi^2$<br>Goodness-of-fit<br>P-value | Occupancy Estimates | |
| --- | --- | --- | --- | --- | --- | --- |
|  |  |  |  |  | Mean | Var. |
| $p$ (CAM + SAV + YR + SITE)<br>$\psi$ (YR) | 826.73 | 0.00 | 0.47 | 0.327 | 0.543 | 0.160 |
| $p$ (CAM + SAV + YR + SITE + TN)<br>$\psi$ (YR) | 828.67 | 1.94 | 0.18 | 0.363 | 0.545 | 0.160 |
| $p$ (CAM + SAV + YR + SITE)<br>$\psi$ (YR + MGMT) | 828.70 | 1.97 | 0.18 | 0.359 | 0.540 | 0.166 |
| $p$ (SAV + YR + SITE)<br>$\psi$ (YR) | 828.73 | 2.00 | 0.17 | 0.353 | 0.548 | 0.153 |

**Table S3:** Sensitivity analysis of species shifts in circular activity distributions, by adjusting the threshold value of human occupancy ± 0.1 from the mean. P-values are given for tests on species shifts using each threshold value. Sig. indicates the observed significance of shifts using the mean threshold value (0.54): + < 0.1, * < 0.05, ** < 0.01, *** < 0.001. The number of significant results (*p-*value < 0.05) using different threshold values is given, for which 3 indicates significance using all thresholds and 0 indicates no significance for any threshold.

| Species | Human occupancy threshold |  |  | Sig. | # sig. |
| --- | --- | --- | --- | --- | --- |
|  | 0.44 | 0.54 | 0.64 |  |  |
| **Apex predators** | 0.035 | 0.019 | 0.016 | * | 3 |
| Hyena | 0.012 | 0.014 | 0.005 | * | 3 |
| Leopard | 0.044 | 0.060 | 0.056 | + | 1 |
| Lion | 0.518 | 0.266 | 0.286 |  | 0 |
| **Ungulates** | 0 | 0 | 0 | *** | 3 |
| Aardvark | 0.001 | 0.012 | 0.017 | * | 3 |
| Buffalo | 0.451 | 0.406 | 0.238 |  | 0 |
| Bushbuck | 0 | 0 | 0 | *** | 3 |
| Duiker | 0.027 | 0.019 | 0.019 | * | 3 |
| Elephant | 0.068 | 0.178 | 0.395 |  | 0 |
| Hartebeest | 0.363 | 0.229 | 0.233 |  | 0 |
| Hippopotamus | 0 | 0 | 0 | *** | 3 |
| Kob | 0 | 0 | 0 | *** | 3 |
| Oribi | 0 | 0 | 0.001 | *** | 3 |
| Reedbuck | 0 | 0 | 0 | *** | 3 |
| Roan Antelope | 0.026 | 0.076 | 0.107 | + | 1 |
| Warthog | 0 | 0 | 0.001 | *** | 3 |
| Waterbuck | 0.053 | 0.118 | 0.098 |  | 0 |

